# Germinal center-derived broadly neutralizing antibodies adapt to SARS-CoV-2 antigenic drift

**DOI:** 10.1101/2022.01.26.477937

**Authors:** Kiyomi Shitaoka, Yohei Kawano, Akifumi Higashiura, Yoko Mizoguchi, Takao Hashiguchi, Norihisa Nishimichi, Shiyu Huang, Ayano Ito, Akima Yamamoto, Shun Ohki, Miyuki Kanda, Tomohiro Taniguchi, Yasuyuki Yokosaki, Satoshi Okada, Takemasa Sakaguchi, Tomoharu Yasuda

**Author notes:** These authors contributed equally to this work (Kiyomi Shitaoka, Yohei Kawano).

## Abstract

The outbreak of SARS-CoV-2 variant Omicron which harbors a striking number of mutations in the spike protein has been raising concerns about the effectiveness of vaccines and antibody treatment^1^. Here, we confirmed a substantial reduction in neutralizing potency against Omicron in all convalescent and vaccinated sera. However, we found that some people infected by the early strain show relatively higher neutralization to Omicron. From those B cells, we developed neutralizing antibodies inhibiting broad variants including Delta and Omicron. Unlike reported antibodies, one had an extremely large interface and widely covered receptor binding motif of spike, thereby interfering with diversified variants. Somatic mutations introduced by long-term germinal center reaction contributed to the key structure of antibodies and the universal interaction with spike variants. Recalling such rare B cells may confer sustainable protection against SARS-CoV-2 variants emerging one after another.

## Main

The antigenic drift of the SARS-CoV-2 RNA virus causes immune evasion through the accumulated mutations in the spike (S) protein that can increase the infectivity of the virus and its transmissibility in the host^2^. The emergence of the SARS-CoV-2 Beta (B.1.351) and Delta (B.1.617) variants raised concern that progress of antigenic drift actually enhanced transmissibility, severity, and mortality of the disease^3^. Recently in November 2021, the Omicron (B.1.1.529) variant was detected and is rapidly spreading worldwide^4^. A striking feature of Omicron is a large number of mutations in the S protein which causes a substantial threat to the efficacy of the current COVID-19 vaccine and antibody therapies^1^. The Omicron variant has as many as 34 mutations in the S protein compared to the original SARS-CoV-2 strain, Wuhan-Hu-1 (**Fig. 1a**). Fifteen of them are clustered in the receptor-binding domain (RBD) which is a primary target of neutralizing antibodies produced after infection or vaccination, including nine mutations located in the receptor-binding motif (RBM), an RBD subdomain that interacts directly with the host receptor ACE2^5^. While recent studies indicated a reduced sensitivity of Omicron to developed therapeutic monoclonal antibodies (mAbs) and COVID-19 convalescent sera^5-8^, little is known about the properties of antibody which can neutralize Omicron. To investigate the mechanism of action and structural basis of broadly neutralizing antibodies from early COVID-19 convalescent individuals will be useful for the development of therapeutic mAbs and vaccines effective to variants emerging one after another. Here we characterized broadly neutralizing mAbs obtained from donors predetermined neutralizing activity against Omicron.

**Fig. 1.**
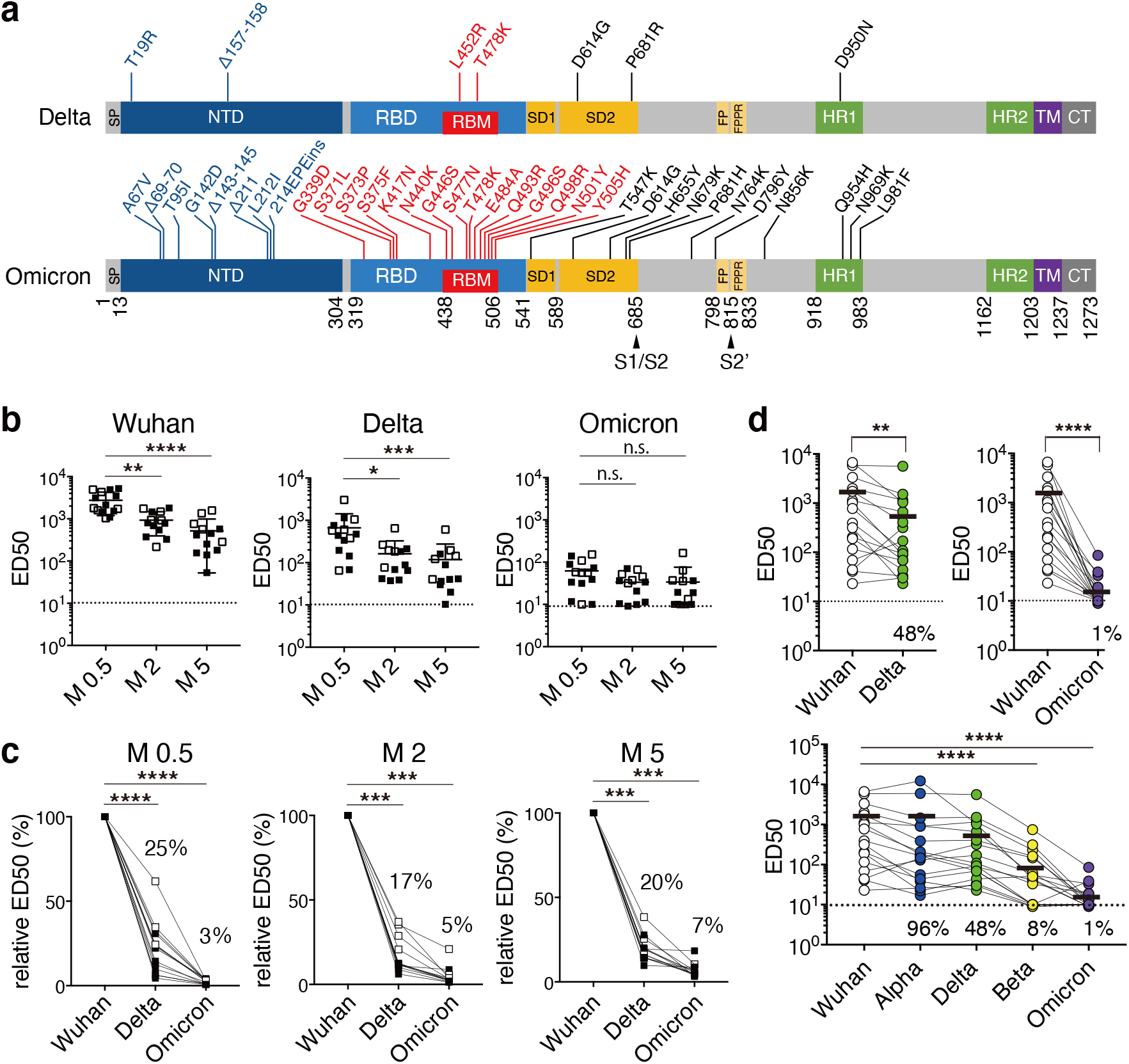
Severe reduction of neutralizing activity against Omicron SARS-CoV-2 strain in sera from COVID-19 convalescent and two-dose vaccinated individuals. **a**, Mutational landscape of Delta and Omicron are shown in the structure of SARS-CoV-2 spike protein with domains and cleavage sites. **b**, Neutralizing activity (ED_50_) against SARS-CoV-2 pseudovirus of Wuhan, Delta, or Omicron strains in sera from individuals, sampled at 0.5 (M0.5), 2 (M2), or 5 months (M5) post-second dose vaccination either from Pfizer (closed squares) or Moderna (open squares). Data are mean±SD (n =14, 13, 13 for M0.5, M2, M5, respectively) from three independent experiments. The dotted line indicates the limit of detection (ED_50_ =10). *p< 0.05, **p< 0.01, ***p< 0.001, ****p< 0.0001, n.s.= not significant. **c**, A pairwise analysis on relative ED_50_ values against SARS-CoV-2 pseudovirus of Delta or Omicron to Wuhan strain at M0.5, M2 or M5. The percentage to Wuhan is shown on each graph. ***p< 0.001.****p< 0.0001. **d**, Pairwise neutralizing antibody titers (ED_50_) of COVID-19 convalescent individuals (n =17) against SARS-CoV-2 pseudovirus of Delta or Omicron to Wuhan strain. **p< 0.01, ****p< 0.0001. The lower panel shows the ED_50_ of each individual against indicated variants. The bold bar indicates the mean from two independent experiments, and the percentages of relative ED_50_ of other variants to Wuhan strain are shown on each graph. The dotted line indicates the limit of detection (ED_50_ =10). A paired two-sided nonparametric test with multiple comparisons of other variants to Wuhan strain was performed. **p< 0.01, ****p< 0.0001.

### Neutralization of Omicron

To investigate the extent of immune evasion of Delta and Omicron variants against vaccine-elicited serum antibodies, we performed a neutralization assay using pseudovirus expressing S protein from Wuhan-Hu-1, Delta (B.1.617.2), or Omicron (B.1.1.529) on the viral membrane. We included fourteen subjects vaccinated twice with BNT162b2 (Pfizer-BioNTech mRNA vaccine, N=8) or mRNA-1273 (Moderna mRNA vaccine, N=6) at different time points, 0.5, 2, and 5 months after the full two-dose vaccination (**Extended Data Table 1**). As recently reported^5-8^, we observed a significant reduction of neutralizing activities against both Wuhan and Delta five months later than the second vaccination (**Fig. 1b**). Fold reduction of neutralization against Wuhan and Delta was 5.3- and 5.6-fold compared to 0.5 months, respectively. In contrast, we did not observe time-dependent reduction against Omicron most likely because neutralization of Omicron was extremely low even soon after the second vaccination. Indeed, neutralization levels against Omicron were near to non-vaccinated/uninfected level at all time points. Therefore, we next calculated changes of neutralization to Wuhan at different time points. At 0.5 to 5 months, ED_50_ against Delta was reduced to 17-25% compared to Wuhan. On the other hand, ED_50_ against Omicron was extremely reduced to 3-7% compared to Wuhan (**Fig. 1c**). We subsequently examined the neutralization ability of sera from convalescent subjects of early D614G strain (B.1.1) sampled at 8-55 days post-diagnosis by PCR (**Extended Data Table 1**). Neutralization of Delta by convalescent sera was slightly reduced (48% to Wuhan). In contrast, convalescent sera extremely reduced neutralization against Omicron in all tested subjects (1% to Wuhan). Next, we further tested sera against additional variants, Alpha (B.1.1.7) and Beta (B.1.351), and found that extent of reduction is largely different in each variant (**Fig. 1d**). Thus, as reported by others, we confirmed a substantial reduction in neutralizing potency against Omicron in all convalescent and vaccinated sera. Not only vaccinated, but most of the convalescent sera also barely neutralized or could not inhibit Omicron pseudovirus. Collectively, these results suggest that individuals fully vaccinated or infected by the early SARS-CoV-2 strain remain at risk for Omicron infection.

### Identification of SARS-CoV-2 broadly neutralizing mAbs

While neutralizing activity elicited by vaccination or natural infection largely fails to inhibit Omicron, we noticed that some convalescent people maintain the neutralizing activity even though at a low level. We conceived that identification of potent broadly neutralizing antibodies from COVID-19 convalescent individuals infected by early D614G strain may be useful for prospective variants, not only Omicron, and provide new insights for ideal therapeutic antibodies. Thus, we next precisely analyzed blood samples from convalescent subjects to establish donor criteria for isolation of broadly neutralizing mAbs. S trimer-specific IgM, IgG, and IgA levels were analyzed in the different periods of hospitalization, 8-10, 11-15, and 17-55 days from PCR diagnosis. S trimer-specific antibody level was the highest in IgG isotype from convalescent people after 17-55 days of hospitalization (**Extended Data Fig. 1a**). Among IgG subtypes, IgG1 was highest compared to IgG2 and IgG3 (**Extended Data Fig. 1b**). Neutralization assay to authentic SARS-CoV-2 indicated that sera from donors of 17-55 days of hospitalization as well as oxygenation treated rather than mild or moderate convalescent donors have the highest neutralization activity. However, we did not observe a clear correlation of neutralizing activities with donor age (**Extended Data Fig. 1c-e**). Therefore, collecting blood from convalescent donors on the day of discharge longer than 17 days of hospitalization with severe symptoms such as oxygenation was thought to be useful criteria to find B cells expressing antibody highly potent in neutralization.

In these criteria, we found two donors (NCV1 and NCV2) showing relatively higher neutralization activity to Omicron (**Fig. 2a**). The clinical characteristics of convalescent subjects including these two patients are summarized in **Extended Data Table 2**. Both patients were highly aged (NCV1 and NCV2 are 93- and 87-year-old, respectively) and hospitalized for 55 days because of severe illness. Peripheral blood mononuclear cells from each donor as well as donors who did not show inhibition of Omicron but were hospitalized for more than 17 days (NCV4, 7, 8) were served for single-cell sorting and processed to produce mAbs (**Extended Data Fig. 2a**). The S trimer binding CD19+IgD-switched live memory B cells were single-cell sorted and we obtained full-length paired IgH and IgL chains with an original IgG subtype and sequence. We recovered 52 sequences from NCV1 and NCV2 together with 50 sequences from control samples (NCV4, 7, and 8) to produce monoclonal IgG antibodies. Among obtained 102 human IgG sequences, IGHV3-30, IGKV1-5, or IGKV3-20 occupied more than 10% each out of whole sequences (**Extended Data Fig. 2b**). These three are previously reported as the predominant V genes used in SARS-CoV-2 S-specific antibodies^9^, indicating that approach allows us to obtain common rearrangements found in potent neutralizing SARS-CoV-2 mAbs from humans. IGHV3-30 and IGKV1-5 were also highly representative in NCV1 and NCV2. Of the 102 IgG mAbs sorted by the S trimer, IgG1, IgG2, and IgG3 subclasses were 86%, 6%, and 8%, respectively. The kappa and lambda chains are comparably used. Thus, IgG1 is dominantly generated against S protein as previously reported^10^ (**Fig. 2b**). Notably, 77% of mAbs acquired somatic mutations from low to high numbers (**Fig. 2c**). The amino acid replacement profile showed accumulated mutations in CDR1 and CDR2 more than FR regions in both VH and VL, indicating that most of S trimer specific antibodies were an outcome of affinity-based selection in the germinal center (GC) (**Fig. 2d**). To find neutralizing mAbs effective to SARS-CoV-2, we carried out antibody neutralization of the authentic SARS-CoV-2 using VeroE6 cells expressing the transmembrane serine protease TMPRSS2. As a result, we identified four mAbs, NCV1SG17, NCV1SG23, NCV2SG48, and NCV2SG53, all of which neutralized D614G authentic SARS-CoV-2 at lower than 1 μg/ml of IC_50_ value. These mAbs were further tested for neutralization against the authentic Delta variant. Three out of four mAbs, NCV1SG17, NCV1SG23, and NCV2SG53 neutralized Delta variant, however, neutralizing activities were reduced by 2.6 to 43-fold. In contrast, NCV2SG48 neutralized Delta at the same level as D614G strain (**Fig. 2e**). Thus, we identified four neutralizing mAbs which are effective to D614G and Delta variants.

**Fig. 2.**
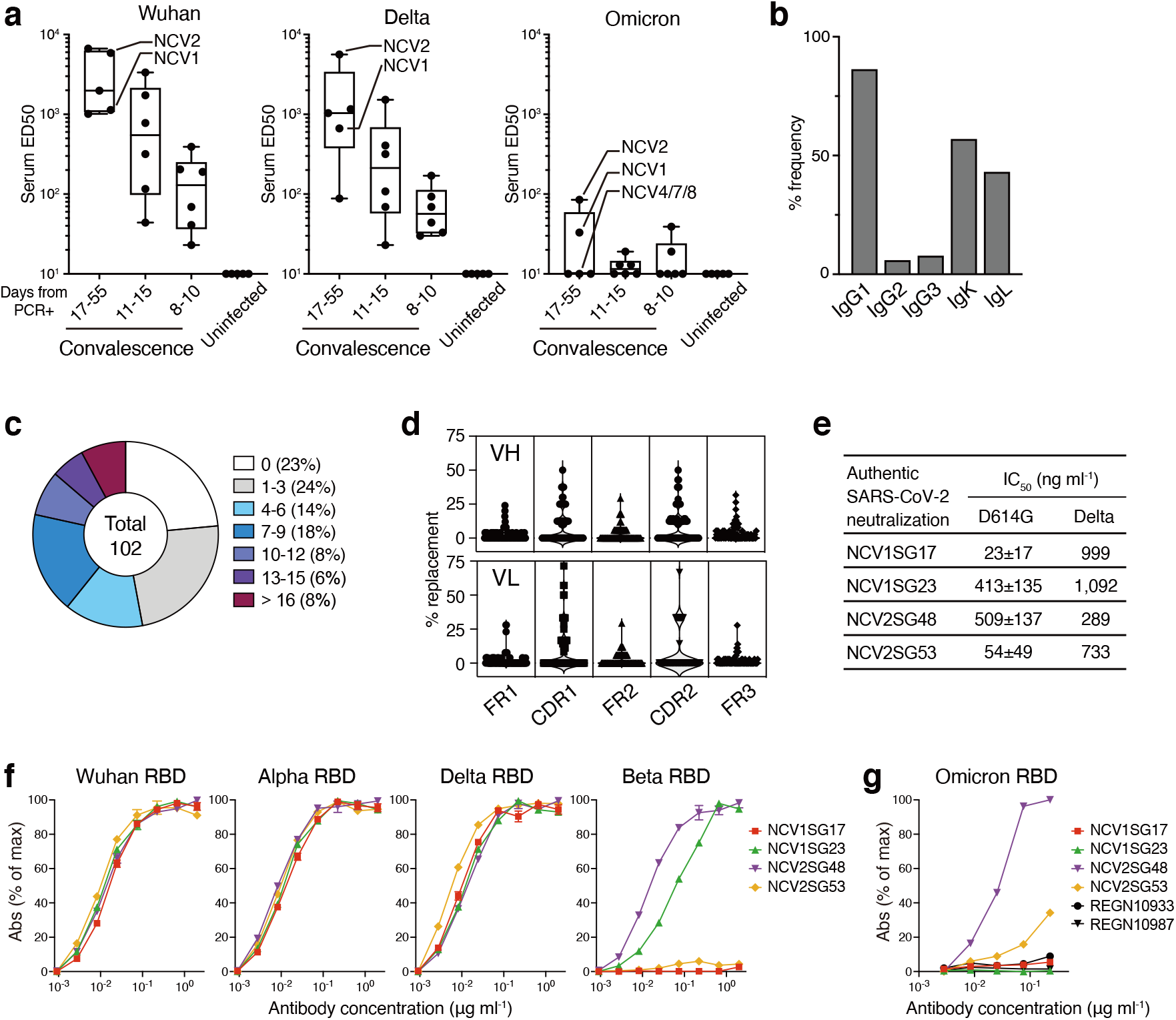
Generation of neutralizing mAbs from convalescence patients showed neutralization to Delta and Omicron. **a**, Serum neutralizing antibody titers of COVID-19 convalescent individuals subgrouped by hospitalization period from PCR result or uninfected healthy donors. ED_50_ values against SARS-CoV-2 pseudovirus of Wuhan, Delta, or Omicron were determined. Subjects used for mAb production were indicated. **b-d**, Sequence of generated 102 mAbs from NCV1, 2, 4, 7, and 8 are analyzed. **b**, Frequency of IgG subclasses and light chain isotypes. **c**, Distribution of VH+VL amino acid mutation numbers. **d**, Percentages of amino acid replacement in FR and CDR region of VH and VL. **e**, The IC_50_ neutralization values of each mAb to indicated authentic SARS-CoV-2 viruses. Mean±SD of three independent results (D614G) or data from a single result (Delta) are shown. **f, g**, Binding properties of indicated mAbs against RBDs of SARS-CoV-2 Wuhan-Hu-1 or its variants.

### Neutralizing properties of identified mAbs

To evaluate whether identified neutralizing mAbs are effective to Omicron and other variants, we first tested mAbs for binding to various RBD proteins because neutralizing antibodies are known to bind RBD to block interaction with ACE2 or viral entry to target cells. As expected, all four mAbs bound specifically with high affinity to monomeric Wuhan-Hu-1 RBD, and that dissociation constant (Kd) was 2.46 to 6.09 nM (**Extended Data Fig. 3**). Next, we evaluated the binding ability of four mAbs, NCV1SG17, NCV1SG23, NCV2SG48, and NCV2SG53, with mutated RBDs. All four mAbs bound to Alpha and Delta at a similar level with Wuhan (**Fig. 2f**). Amino acid substitutions of RBD in SARS-CoV-2 variants used in this study are summarized in **Extended Data Fig. 4a**. We also included variants acquired point mutation only at K417, L452, or E484, which are known to decrease sensitivity to antibody neutralization^11,12^. RBD binding of all four mAbs was not affected by those mutations (**Extended Data Fig. 4b**). Of note, NCV1SG23 and NCV2SG48 mAbs showed binding to all variants including, Alpha, Beta, Kappa, Delta, Delta plus, Lambda, and Omicron, except for NCV1SG23 lost binding to Omicron (**Fig. 2f, g and Extended Data Fig. 4b**). As recently reported, REGN10933 and REGN10987 mAbs, also known as Casirivimab and Imdevimab, respectively, lost binding to Omicron (**Fig. 2g**) ^5-7^.

To confirm the neutralizing ability of NCV1SG17, NCV1SG23, NCV2SG48, and NCV2SG53 against SARS-CoV-2 variants, we carried out pseudovirus neutralization assay corresponding to Wuhan-Hu-1, D614G, Alpha, Beta, Kappa, Delta, Delta plus, Lambda, and Omicron. As expected from the above results, NCV2SG48 neutralized all variants including Omicron (**Fig. 3a and Extended Data Fig. 5a**). Although NCV2SG48 showed 13-fold decrease in neutralization activity against Omicron compared to parental D614G, 798 ng/ml of IC_50_ value to Omicron is equivalent level with reported neutralizing antibody Sotrovimab, which is derived from S309 mAb isolated from SARS-CoV infected individual, inhibiting Omicron by 260-917 ng/ml of IC_50_ value^5,6^. While NCV1SG17, NCV1SG23, and NCV2SG53 did not show neutralization of Omicron, those mAbs neutralized most of the other variants. Upon fact that IC_50_ values of NCV2SG53 are 10-fold higher than those of NCV2SG48 against Delta and some other variants, we tested an antibody cocktail consisting of NCV2SG48 and NCV2SG53. These antibodies in the cocktail did not obstruct each other and broadly neutralized all variants (**Fig. 3b, c and Extended Data Fig. 5b, c**). It has been demonstrated that the use of mAbs targeting the S protein is a powerful way to treat COVID-19 patients, however, the emergence of antibody-resistant escape mutants still remains as concern^13,14^. To evaluate the susceptibility of our neutralizing antibodies to the appearance of escape mutants, we cultured authentic SARS-CoV-2 virus in the presence of each mAb. We detected multiple escape mutants from the culture with NCV1SG17 or NCV2SG53 and identified S494P mutation or E484D/G485D/G485R mutations, respectively. However, we detected none of the escape mutants from seven independent cultures under the presence of NCV1SG23 or NCV2SG48 alone suggesting that these two mAbs are highly resistant to spontaneous mutations (**Extended Data Table 3**). Collectively, the NCV2SG48+NCV2SG53 antibody cocktail will be attractive for the treatment of COVID-19 patients infected by SARS-CoV-2 variants.

**Fig. 3.**
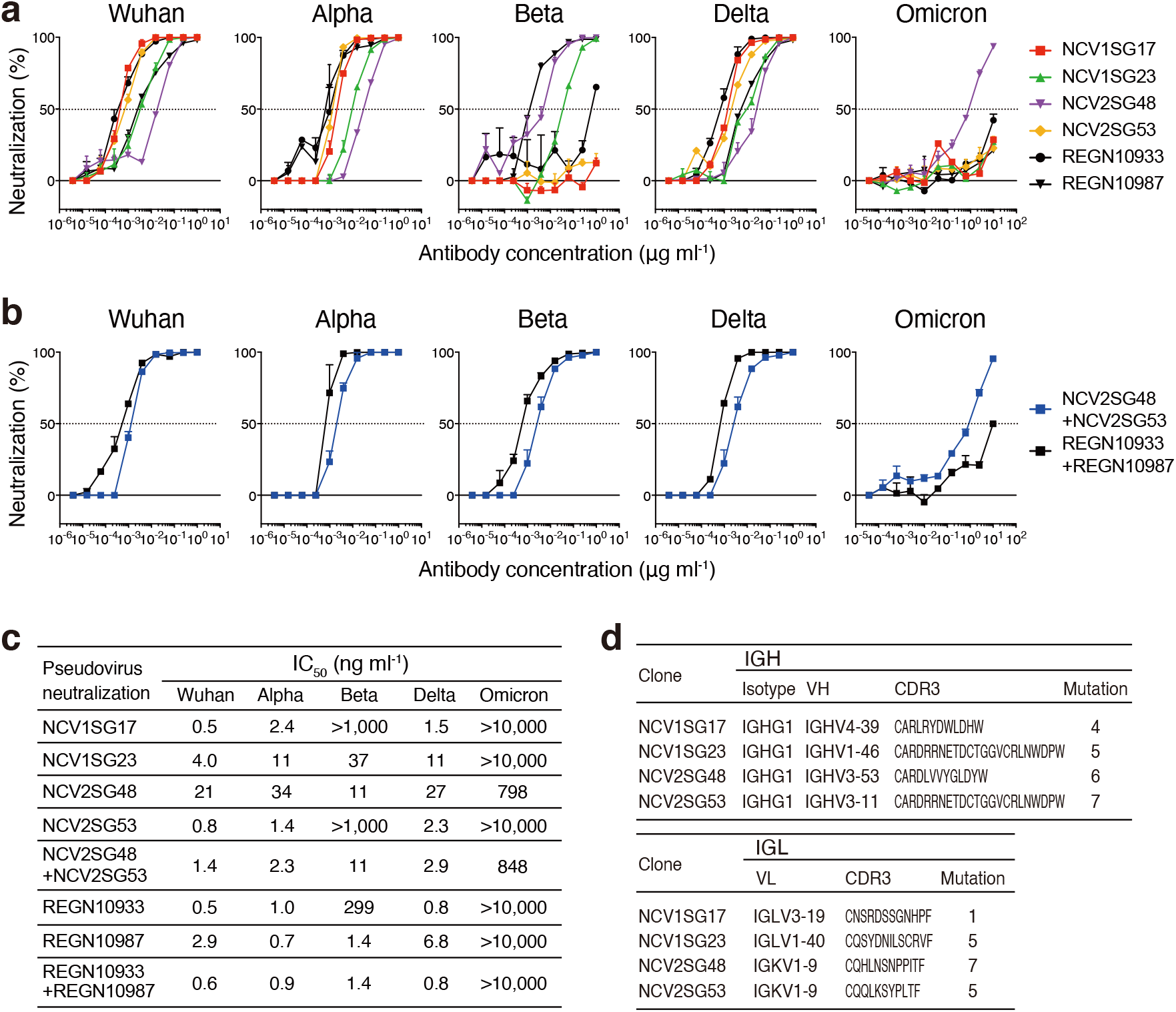
Neutralizing potency of mAbs against SARS-CoV-2 Delta and Omicron. **a**, Dose-response analysis of the neutralization by each mAb NCV1SG17, NCV1SG23, NCV2SG48, NCV2SG53, REGN10933 and REGN10987 on indicated variants of SARS-CoV-2 pseudovirus. The horizontal dotted line on each graph indicates 50% neutralization. Data are mean±SEM of technical duplicates from 2 to 3 independent experiments. **b**, Dose-response analysis of the neutralization by the mixture of mAbs NCV2SG48 and NCV2SG53, or REGN10933, and REGN10987 on indicated variants of SARS-CoV-2 pseudovirus. The horizontal dotted line on each graph indicates 50% neutralization. Data are mean±SEM of technical duplicates from 2 to 3 independent experiments. **c**, The IC_50_ of each antibody and the mixture of mAbs to indicated SARS-CoV-2 variants. **d**, VH, CH, and VL usage, CDR3 sequence, and the number of amino acid mutations of neutralizing mAbs.

### Structural basis of broadly neutralizing mAbs

To gain insight into how NCV2SG48 and NCV2SG53 neutralize broad variants, we determined the X-ray crystal structure of antigen-binding fragments (Fabs) in complex with the RBD. From the structure analysis, we found that both antibodies differently recognize receptor binding module (RBM) to inhibit ACE2-binding close to each other (**Fig. 4a**). Neutralizing antibodies of SARS-CoV-2 can be classified based on binding region^6,15^. We found that NCV2SG48 is a typical class 1 antibody, while NCV2SG53 recognizes an intermediate region between class 1 and class 2 (**Extended Data Fig. 6, 7a**). Notably, NCV2SG48 interacted with RBD by extensive large interface covering almost the entire ACE2-binding region and the adjacent area, which accounts for 133-203% compared to the interface of REGN10933, REGN10987, or S309 (**Extended Data Fig. 7b-d**). NCV2SG48 could form 33 hydrogen bonds including water-mediated bonds which possibly accept even bulky amino acids that emerged in Omicron (**Extended Data Fig. 7e**). Structure analysis suggests that somatic mutation generated five additional hydrogen bonds which contributed to cover an extended area on RBM (**Extended Data Fig. 8a**). In contrast, NCV2SG53 is highly dependent on interaction with E484 on RBD by forming 14 hydrogen bonds around E484. Somatic mutation of T57S in HC-CDR2 could contribute to reduce the steric hindrance with F490 of RBD and contributed to the higher affinity around E484 (**Extended Data Fig. 8b**). This explains why E484 mutation emerged in Beta and Omicron abolished neutralization by NCV2SG53. The importance of E484 is further confirmed by escape mutant analysis (**Extended Data Table 3**). Collectively, NCV2SG53 can neutralize diversified variants that do not have E484 mutation and NCV2SG48 is highly resistant to mutations by covering an extensive wide area on RBM. In both cases, somatic mutation introduced via GC reaction is thought to play a critical role in universal inhibition against diversified variants.

**Fig. 4.**
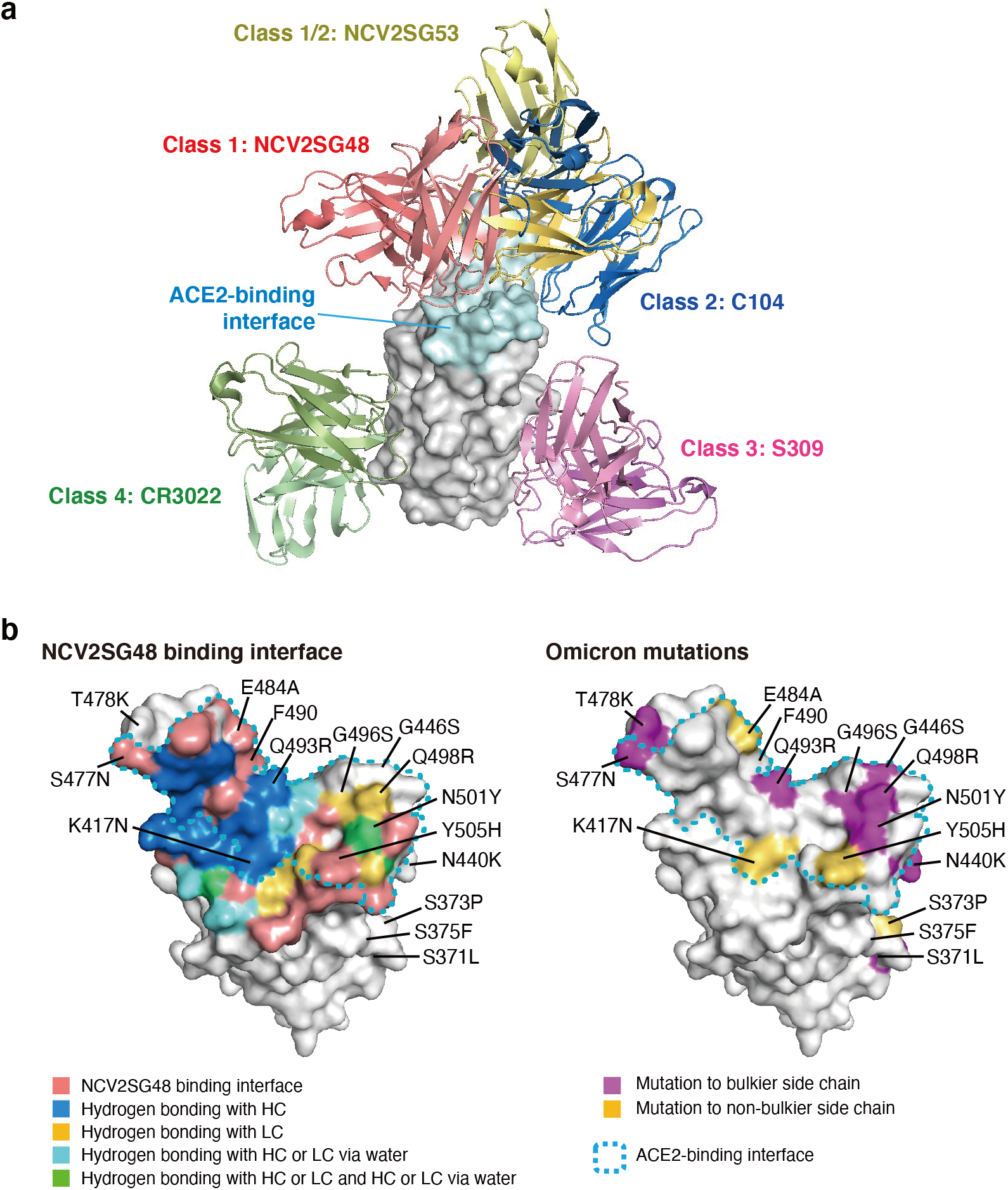
Structure and epitope map of NCV2SG48 and NCV2SG53 binding with the SARS-CoV-2 RBD. **a**, The overview of Fab variable regions of Class 1: NCV2SG48 (*light red*), Class 1/2: NCV2SG53 (*yellow*), Class 2: C104 (*blue*; PDB ID:7K8U), Class 3: S309 (*magenta*; PDB ID:7R6W) and Class4: CR3022 (*green*; PDB ID:6W41) complexed with RBD. RBD is shown as a *gray* surface model and the ACE2-binding interface is colored in *cyan*. NCV2SG53 binds to the intermediate epitope region between class 1 and class 2. **b**, *Left*: Antibody (NCV2SG48) binding sites are shown on the surface model of RBD. NCV2SG48 binding interface is shown in *light red* and the ACE2-binding site is surrounded by a *cyan* dotted line. Residues of hydrogen bonded to HC (*blue*), LC (*yellow*), HC via water molecule (*cyan*) are shown on the RBD surface model. Residues from two hydrogen bonds with Fab are shown in *green. Right*: Mutated residues of omicron variant on RBD surface model. Mutation to bulkier side chains and to a not bulkier side chains are shown in *purple* and *yellow*.

## Discussion

We aimed to address how antibodies produced in convalescent or vaccinated individuals by early strain neutralize a broad variant including heavily mutated Omicron. Here we found that both convalescent and vaccinated individuals elicited polyclonal antibodies to early viral sequence barely possessed antibodies inhibiting Omicron. Based on our results, the frequency of broadly neutralizing antibodies in circulating B cells was around 4% (4 out of 102 mAbs) among S trimer-specific B cells recovered from convalescent patients with D614G strain but only 1% was effective to Omicron. Neutralization assay with sera against variants was useful to identify blood donors to isolate broadly neutralizing mAbs in addition to donor criteria, over 17 days of hospitalization with severe illness. NCV2SG48 antibody was rearranged by IGHV3-53 and IGKV1-9 (**Fig. 3d**), which combination was not detected in mAbs other than NCV2SG48 suggesting broadly neutralizing antibodies may be derived from minor responders. This may explain why immunized sera show extremely reduced neutralizing titers to Omicron. Based on the fact that significantly higher neutralization activity was detected only in donors who experienced long-term hospitalization, the accumulated somatic mutation in the antibody must play important role in increasing neutralization activity and the generation of broadly neutralizing antibodies. To our surprise, we could isolate broadly neutralizing mAbs from 87- and 93-year-old donors even though it is accepted that elderly people are at high risk for serious symptoms or death^16-19^. It is interesting whether the ability to generate neutralizing antibodies correlates with clinical outcomes in elderly people. Our antibody cocktail consisting of NCV2SG48 and NCV2SG53 can be useful for the treatment of COVID-19 patients infected by future variants.

Hybrid immunity brought by sequential immunization “infection-then-vaccination” is highly effective to polymutant spike or diverse sarbecovirus spike proteins^20-22^. The presence of universal neutralizing antibodies in early COVID-19 convalescent individuals suggests that recalling such rare but exceptionally flexible B cells may contribute to hybrid immunity and confer sustainable protection against SARS-CoV-2 pandemics. This might be achieved if convalescent or vaccinated individuals with the original strain are boosted by Omicron-specific vaccine which may become available in the near future. Our results suggest that the human immune system barely adapts to highly extended antigenic drift if only repeating vaccine with the original sequence but may overcome this issue by the “original-then-variant” cross vaccination based on the GC-mediated adaptation to SARS-CoV-2 antigenic drift.

## Methods

### Convalescent or vaccinated human donors

Volunteers aged 23 to 93 with a history of convalescent COVID-19 were enrolled from April 2020 to January 2021. Blood samples were collected on the day or one day before discharging from the hospital after symptom resolution. Duration is the time between PCR positive and blood sample collection. Healthy volunteers aged 22 to 62 were enrolled and the blood samples were collected before or after vaccination with Pfizer-BioNTech Comirnaty or Moderna mRNA-1273 COVID-19 vaccine. Detailed information on the cohort is in **Extended Data Table 1**. PBMCs and plasma samples were isolated by density gradient centrifugation with Ficoll-Paque PLUS (GE Healthcare) and stored at -80°C until use.

### Flow cytometry for single-cell sorting

For single-cell sorting, PBMCs were treated with FcX blocking antibodies (BioLegend, #4422302) to reduce non-specific labeling of the cells. PBMCs were stained with S trimer-Strep-tag, CD19-APC-Cy7 (BioLegend, #302217), and IgD-FITC (BioLegend, #348206) for 20 min on ice. After washing, cells were stained with Strep-Tactin XT-DY649 (IBA) for 20 min on ice. The cells were resuspended in FACS buffer (PBS containing 1% FCS, 1 mM EDTA, and 0.05% NaN3) supplemented with 0.2 μg/ml propidium iodide (PI) to exclude dead cells. Cell sorting was performed on FACSAria II (BD Biosciences) to isolate S trimer^+^ CD19^+^ IgD^-^ cells from the PI^-^ live cell gate. Cells were directly sorted into a 96-well PCR plate. Plates containing single-cells were stored at -80°C until proceeding to RT-PCR.

### Single-cell RT-PCR and monoclonal antibody production

Single-cell sorted PCR plates were added by 2 μl of pre-RT-PCR mix containing the custom reverse primers to each well. After heating at 65°C for 5 min, plates were immediately cooled on ice. 2 μl of the pre-RT-PCR2 (PrimeScript™ II Reverse Transcriptase, Takara Bio) mix was added to each well. For RT reaction, samples were incubated at 45°C for 40 min followed by heating at 72°C for 15 min, then cooled on ice. For PCR amplification of full-length immunoglobulin heavy and light chain genes, PrimeSTAR DNA polymerase (Takara Bio) and custom primers were used. For first PCR, the initial denaturation at 98°C for 1 min was followed by 25 cycles of sequential reaction of 98°C for 10 sec, 55°C for 5 sec, and 72°C for 1.5 min. For second PCR, the initial denaturation at 98°C for 1 min was followed by 35 cycles of sequential reaction of 98°C for 10 sec, 58°C for 5 sec, and 72°C for 1.5 min. PCR fragments were assembled into a linearized pcDNA vector using NEBuilder HiFi DNA Assembly Master Mix (New England Biolabs) according to the manufacturer’s instructions. The pcDNA3 (Invitrogen) vectors containing an Ig light chain gene and the pcDNA4 (Invitrogen) vectors containing an Ig heavy chain gene were simultaneously transfected into Expi293 cells using Expi293 Expression System Kit (Thermo Fisher Scientific). Four days after the transfection, the culture supernatants were collected and subjected to ELISA.

### Production of recombinant S trimer and RBD

Production of recombinant S trimer was described previously^23^. Briefly, the human codon-optimized nucleotide sequence encoding for the S protein of SARS-CoV-2 (GenBank: MN994467) was synthesized commercially (Eurofins Genomics). A soluble version of the S protein (amino acids 1–1213), including the T4 foldon trimerization domain, a histidine tag, and a strep-tag, was cloned into the mammalian expression vector pCMV. The protein sequence was modified to remove the polybasic cleavage site (RRAR to A), and two stabilizing mutations were also introduced (K986P and V987P; wild-type numbering)^24^. The gene encoding RBD of SARS-CoV-2, Wuhan-Hu-1, was synthesized and cloned into vector pcDNA containing a human Ig leader sequence and C-terminal 6xHis tag. RBD mutants were generated by overlap PCR using primers containing mutations. The vector was transfected into Expi293 cells and incubated at 37°C for 4 days. Supernatants were purified using Capturem™ His-Tagged Purification kit (Takara Bio), then dialyzed by PBS buffer overnight. Protein purity was confirmed by SDS-PAGE. Protein concentration was determined spectrophotometrically at 280 nm.

### ELISA for SARS-CoV-2 antigens

MaxiSorp ELISA plates (Thermo Fisher Scientific) were coated with 2 μg/ml purified RBD or S protein in 1xBBS (140 mM NaCl, 172 mM H3BO3, 28 mM NaOH) overnight at 4°C, and then blocked with blocking buffer containing 1% BSA in PBS for 1 hour. Antibodies diluted in Reagent Diluent (0.1% BSA, 0.05% Tween in Tris-buffered Saline) were added and incubated for 2 hours. HRP-conjugated antibodies were added and incubated for 2 hours. Wells were reacted with the TMB substrate (KPL) and the reaction was stopped using 1 M HCl. The absorbance at 450 nm was measured using iMark microplate absorbance reader (Bio-Rad).

### Affinity measurement using biolayer interferometry (BLI)

The binding affinity of obtained antibodies to RBD was examined by the Blitz system (Sartorius Japan) using protein A-coated biosensors. 10 μg/ml of antibody was captured by the biosensor and equilibrated, followed by sequential binding of each concentration of RBD. For dissociation, biosensors were dipped in PBST for 900 sec. Results were analyzed by Blitz system software.

### Reagents

psPAX2 (Addgene, no.12260) was a gift from Didier Trono. pCDNA3.3_CoV2_B.1.1.7 (Addgene, no.170451) for Alpha-S and pcDNA3.3-SARS2-B.1.617.2 (Addgene, no.172320) for Delta-S proteins, were gifts from David Nemazee^25^. pTwist-SARS-CoV-2 Δ18 B.1.351v1 (Addgene, no.169462) for Beta-S protein was a gift from Alejandro Balazs^26^. Lentiviral vector, pWPI-ffLuc-P2A-EGFP for luciferase reporter assay and pTRC2puro-ACE2-P2A-TMPRSS2 for the generation of 293T cell line susceptible to SARS-CoV-2 infection was created from pWPI-IRES-Puro-Ak-ACE2-TMPRSS2, a gift from Sonja Best (Addgene, no.154987) by In-Fusion® technology (Takara Bio). pcDNA3.4 expression plasmids encoding SARS-CoV-2 S proteins with human codon optimization and 19 a.a deletion of C-terminus (C-del19) from Wuhan, D614G, and Omicron were generated by assembly of PCR products, annealed oligonucleotides, or artificial synthetic gene fragments (Integrated DNA Technologies, IDT) using In-Fusion® technology. For Delta plus, Kappa and Lambda variants, S proteins with only RBD, D614, and P681 mutations were created from pcDNA3.4 encoding human codon-optimized Wuhan S protein (C-del19). LentiX-293T cells (Takara Bio) and 293T cells were maintained in culture with Dulbecco’s Modified Eagle’s Medium (DMEM) containing 10% fetal bovine serum (FBS), penicillin-streptomycin (Nacalai tesque), and 25 mM HEPES (Nacalai tesque).

### Generation of 293T cells stably expressing human ACE2 and TMPRSS2

To generate stable 293T-ACE2.TMPRSS2 cells (293T/TRCAT), lentiviral vector VSV-G-pseudotyped lentivirus carrying ACE2 and TMPRSS2 genes were produced in LentiX-293T cells by transfecting with pTRC2puro-ACE2-P2A-TMPRSS2, psPAX2 (gag-pol), and pMD2G-VSV-G (envelope) using PEI-MAX (Polysciences). Packaged lentivirus was used to transduce 293T cells in the presence of 5μg/mL polybrene. At 72 hours post-infection, the resulting bulk transduced population positive for Human ACE2 expression stained by FITC-anti-ACE2 Antibody (Sinobiological) was sorted by FACS Aria SORP (BD) and maintained in the culture medium in the presence of 2 μg/ml of puromycin.

### Pseudovirus production and neutralization

Pseudoviruses bearing SARS-Cov2 S-glycoprotein and carrying a firefly luciferase (ffLuc) reporter gene were produced in LentiX-293T cells by transfecting with pWPI-ffLuc-P2A-EGFP, psPAX2, and either of S variant from Wuhan, D614G, Alpha, Beta, Delta, Delta plus, Kappa, Lambda, or Omicron using PEI-MAX (Polyscience). Pseudovirus supernatants were collected approximately 72 hours post-transfection and used immediately or stored at -80°C. Pseudovirus titers were measured by infecting 293T/TRCAT cells for 72 hours before measuring luciferase activity (ONE-Glo™ Luciferase Assay System, Promega, Madison, WI). Pseudovirus titers were expressed as relative luminescence units per milliliter of pseudovirus supernatants (RLU/ml). For neutralization assay, pseudoviruses with titers of 1-4×106 RLU/ml were incubated with antibodies or sera for 0.5 hours at 37°C. Pseudovirus and antibody mixtures (50 μl) were then inoculated with 5μg/ml of polybrene onto 96-well plates that were seeded with 50μl of 1 × 104 293T/TRCAT cells /well one day before infection. Pseudovirus infectivity was scored 72 hours later for luciferase activity. The serum dilution or antibody concentration causing a 50% reduction of RLU compared to control (ED_50_ or IC_50_, respectively) were reported as the neutralizing antibody titers. ED_50_ or IC_50_ were calculated using a nonlinear regression curve fit (GraphPad Prism software Inc., La Jolla, CA).

### Neutralization assay with authentic SARS-CoV-2 viruses

VeroE6/TMPRSS2 cells (African green monkey kidney-derived cells expressing human TMPRSS2, purchased from the Japanese Collection of Research Bioresources (JCRB) Cell Bank, JCRB1819) were maintained in DMEM containing 10% FBS and 1 mg/mL G418 at 37 °C in 5% CO2. The SARS-CoV-2/JP/Hiroshima-46059T/2020 strain (B.1.1, GISAID: EPI_ISL_6289932, GenBank accession number: MZ853926), which was isolated from a cluster infection in Hiroshima^27^, was propagated in VeroE6/TMPRSS2 cells and used as test virus. The virus titer was determined by the standard 50% tissue culture infectious dose (TCID50) method and expressed as TCID50/ml as described previously^28^. The serially diluted antibody (50 μl) was mixed with 100 TCID50/50 μl of the virus and reacted at 37°C for 1 hour, then inoculated into VeroE6/TMPRSS2 cells to determine the minimum inhibitory concentration (MIC) or 50% effective dose (ED_50_) of the antibody. SARS-CoV-2 infection was performed in the BSL3 facility of Hiroshima University.

### Escaping SARS-CoV-2 mutants from antibodies

SARS-CoV-2 was incubated with mAb at twice the concentration of the EC50 corresponding to that viral load at 37°C for 60 minutes. After the incubation, 100 μl of the mixture was added to one well of a 24-well plate with confluent VeroE6/TMPRSS2 cells and incubated for 72 hours at 37 °C with 5% CO2. The supernatants were collected as an escape-mutant virus when CPE was manifested. A no-antibody-control was included to confirm the amount of test virus required.

### Virus RNA Sequencing

Viral RNA was extracted from virus-infected culture medium by using Maxwell RSC Instrument (Promega, AS4500). cDNA preparation and amplification were done in accordance with protocols published by the ARTIC network (https://artic.network/ncov-2019) using V4 version of the ARTIC primer set from Integrated DNA Technologies to create tiled amplicons across the virus genome. The sequencing library was prepared using the NEB Next Ultra II DNA Library Prep Kit for Illumina (New England Biolabs, E7645). Paired-end, 300 bp sequencing was performed using MiSeq (Illumina) with the MiSeq reagent kit v3 (Illumina, MS-102-3003). Consensus sequences were obtained by using the DRAGEN COVID lineage software (Illumina, ver. 3.5.6). Variant calling and annotation were performed using the Nextclade website (https://clades.nextstrain.org).

### Preparation of RBD and Fab complexes

RBD and Fab fragment from NCV2SG48 with 6xHis-tag expressed in Expi293F cells (Thermo Fisher Scientific) were purified using Ni-NTA Agarose resin (QIAGEN). Fab fragment of NCV2SG53 was isolated from papain digests of the monoclonal antibody expressed in Expi293F cells using HP Protein G column (Cytiva). Purified each Fab fragments and RBD were mixed in the molar ration of 1:1.2 and incubated on ice for 1 hr. The mixture was loaded onto a SuperdexTM 200 increase 10/300 GL column (Cytiva) equilibrated in 20 mM Tris-HCl pH7.5, 150 mM NaCl for removing the excess RBD. Fractions containing RBD and each Fab were collected and concentrated for crystallization.

### Crystallization, X-ray diffraction data collection, and processing

Crystallization was carried out by the sitting-drop vapor diffusion method at 20 °C. Crystals of RBD-Fab (NCV2SG48) were grown in 2 μl drops containing a 1:1 (v/v) mixture of 7.5 mg ml-1 RBD solution and 0.1 M Bis-Tris pH5.5, 0.5 M ammonium sulfate, 19% PEG3350. Crystals of RBD-Fab (NCV2SG53) were grown in 2 μl drops containing a 1:1 (v/v) mixture of 7.5 mg ml-1 RBD solution and 0.1 M MES pH6.0, 0.25 M ammonium sulfate, 22.5% PEG3350. The single crystals suitable for X-ray experiments were obtained in a few days. X-ray diffraction data collections were performed using synchrotron radiation at SPring-8 beamline BL44XU^29^ in a nitrogen vapor stream at 90 K. The data sets were indexed and integrated using the XDS package^30^, scaled, and merged using the program *Aimless*^31^. The scaling statistics were shown in **Extended Data Table 4**.

### Structure determination and analyses

Phase determinations were carried out by the molecular replacement method using the program Phaser in the PHENIX package^32^ and the program Molrep^33^ with the combination between RBD structure (PDB ID:7EAM) and Fab structures (PDB ID:7CHB and 7CHP) as search models. The structure refinement was performed using the program phenix.refine^34^ and the program coot^35^. The final refinement statistics were shown in **Extended Data Table 4**. Interactions between RBD and Fabs were analyzed using the program PISA^36^. All figures of structures were generated by the program pymol (The PyMOL Molecular Graphics System, Version 1.2r3pre, Schrödinger, LLC.). Class 2/C104 (PDB ID:7K8U), Class 3/S309 (PDB ID:7R6W), and Class4/CR3022 (PDB ID:6W41) open data were used in **Fig. 4**.

### Quantification and statistical analysis

Ordinary One-way ANOVA, Two-way ANOVA, Kruskal-Wallis test, Wilcoxon rank test, and Friedman test were used to compare data. P-value 0.05 was considered statistically significant. Statistical analyses were all performed using GraphPad Prism 9.0 (La Jolla, CA, USA).

## Data and availability

The structure of SARS-CoV-2 RBD in complex with NCV2SG48 and NCV2SG53 Fabs has been deposited in the Protein Data Bank as the PDB ID:7WNB and 7WN2, respectively.

## Acknowledgements

We thank all study participants for our research; Blood donors, medical staff of the Hiroshima University Hospital, Shobara Redcross Hospital, Hiroshima Prefectural Hospital. We thank T. Utsumi, S. Eto, Y. Tamura, T. Kawahara for the arrangement of donors and collecting blood; T. Kawaguchi, N. Tani, and Y. Hayashi for technical assistance; N. Kikkawa for administrative assistance. All lab members for useful discussion and comments. We also thank the staff of the Analysis Center of Life Science, Hiroshima University for the use of their facilities. This work was performed using a synchrotron beamline BL44XU at SPring-8 under the Collaborative Research Program of the Institute for Protein Research, Osaka University. Diffraction data were collected at the Osaka University beamline BL44XU at SPring-8 (Harima, Japan) (Proposal No. 2021B6651). This work was supported by the JSPS KAKENHI Grant Numbers JP17H06937, JP18H02669, JP19K22538, and JP21H02751 to T.Y.; Sumitomo Mitsui Trust Bank-New Corona Vaccine and Therapeutics Development Donation Account to T.S. and T.Y.; Japan Agency for Medical Research and Development (AMED) Research Grant for COVID-19, JP20fk0108453 to T.S. and T.Y; Hiroshima Prefecture-Hiroshima University Government-Academia Collaboration COVID-19 Research Fund to T.S.; AMED Practical Research Project for Rare/Intractable Diseases, JP20fk0108531 to S.O.; AMED Translational Research Program, PreB 20334760 to N.N. and Y.Y.; Ministry of Education, Culture, Sports, Science and Technology grant 20H05773 to T.H.; and JST CREST Grant Number JPMJCR20H8 to T.H..

## Author contributions

K.S., Y.K. Y.M. and T.Y. designed the experiments. T.T. provided blood from patients. K.S., A.H., T.H., and A.Y. produced and purified recombinant proteins. K.S. and N.N. sequenced and produced antibodies. K.S., N.N. and S.H. performed protein binding assays. Y.K. and A.I. performed pseudovirus neutralization assays. T.S. and M.K. performed neutralization and escape assays with authentic SARS-CoV-2. A.H. and A.Y. performed X-ray crystallography and analyzed the structure data. S.Oh. performed bioinformatic analysis. T.H. provided key reagents. K.S., Y.K. and T.Y. prepared the manuscript with input from all authors. Y.Y., S.Ok, T.S., and T.Y. supervised the study.

---

**Extended Data Fig. 1.**
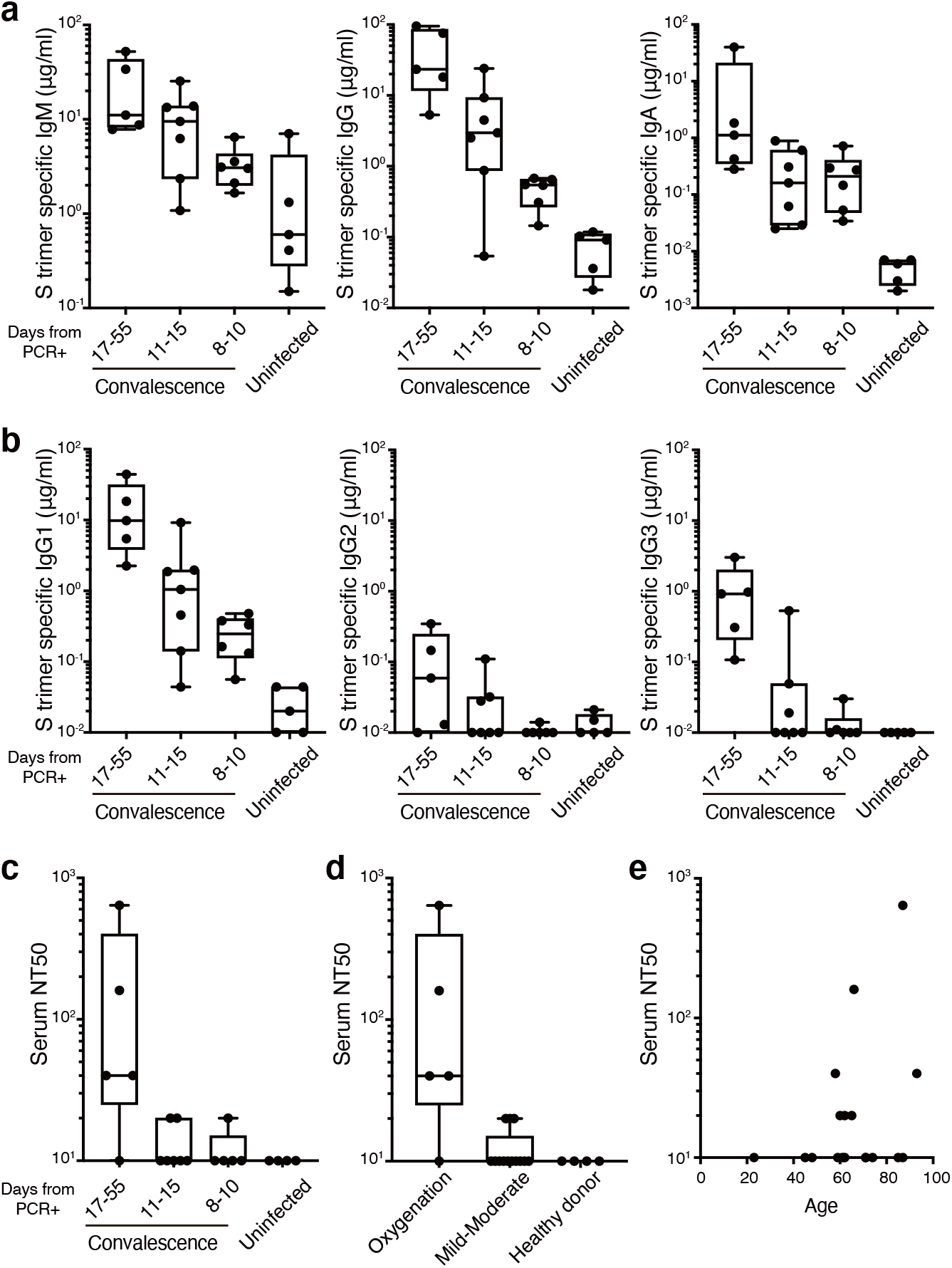
COVID-19 convalescent individuals after the long hospitalization show higher S-specific antibodies and neutralization activity against SARS-CoV-2. **a**, The S timer-specific serum IgM, IgG, and IgA levels, **b**, the S timer-specific serum IgG1, IgG2, and IgG3 levels, and **c**, serum neutralizing antibody titers of COVID-19 convalescent individuals subgrouped by hospitalization period from PCR result or uninfected healthy donors. The neutralizing activity of serum antibodies was evaluated by testing the blocking effect of authentic SARS-CoV-2 D614G virus infection to Vero cells. The 50% neutralization titer (NT_50_) was determined using the half-maximal inhibitory concentration values. **d**, Serum neutralizing antibody titers of COVID-19 convalescent individuals subgrouped by the severity of the disease. **e**, Serum neutralizing antibody titer was plotted to the age of COVID-19 convalescent individuals.

**Extended Data Fig. 2.**
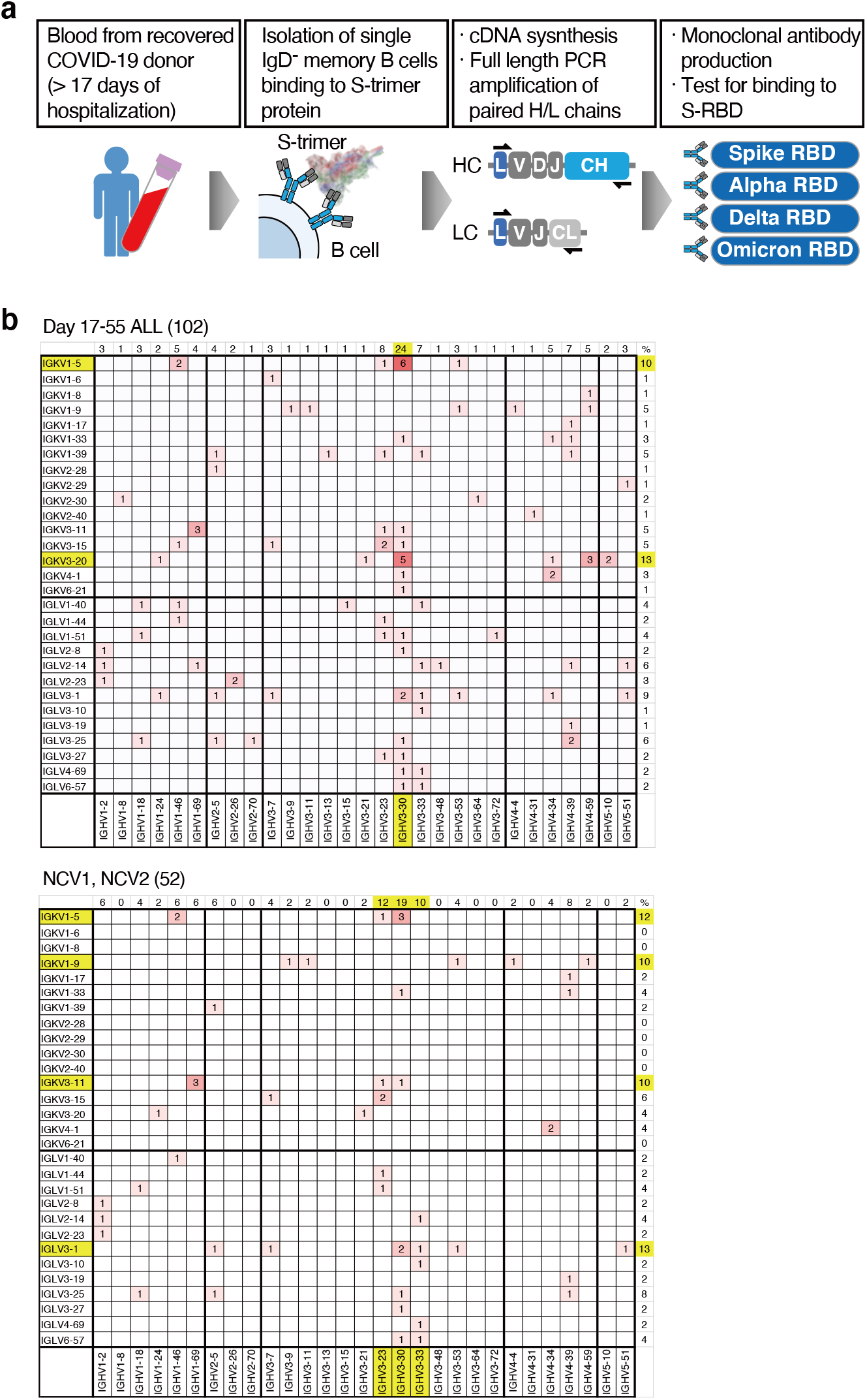
Strategy for S trimer-specific mAb generation and variable gene usage of generated mAbs. **a**, Schema for isolation of S trimer-specific mAbs from single-cell sorted Ig-switched B cells in the blood of COVID-19 convalescent individuals. **b**, The variable (V) gene frequencies for paired heavy (*x-axes*) and light (*y-axes*) chains of isolated S trimer-specific mAbs. Numbers of identified V genes in 102 mAbs from 5 donors of day 17-55 hospitalization period (*upper*) or 52 mAbs from NCV1 and NCV2 (*bottom*) are summarized with red color indication. Percent frequency of V genes is shown on the top and right of each panel for VH and VL, respectively. V genes account for over 10% are highlighted by *yellow*.

**Extended Data Fig. 3.**
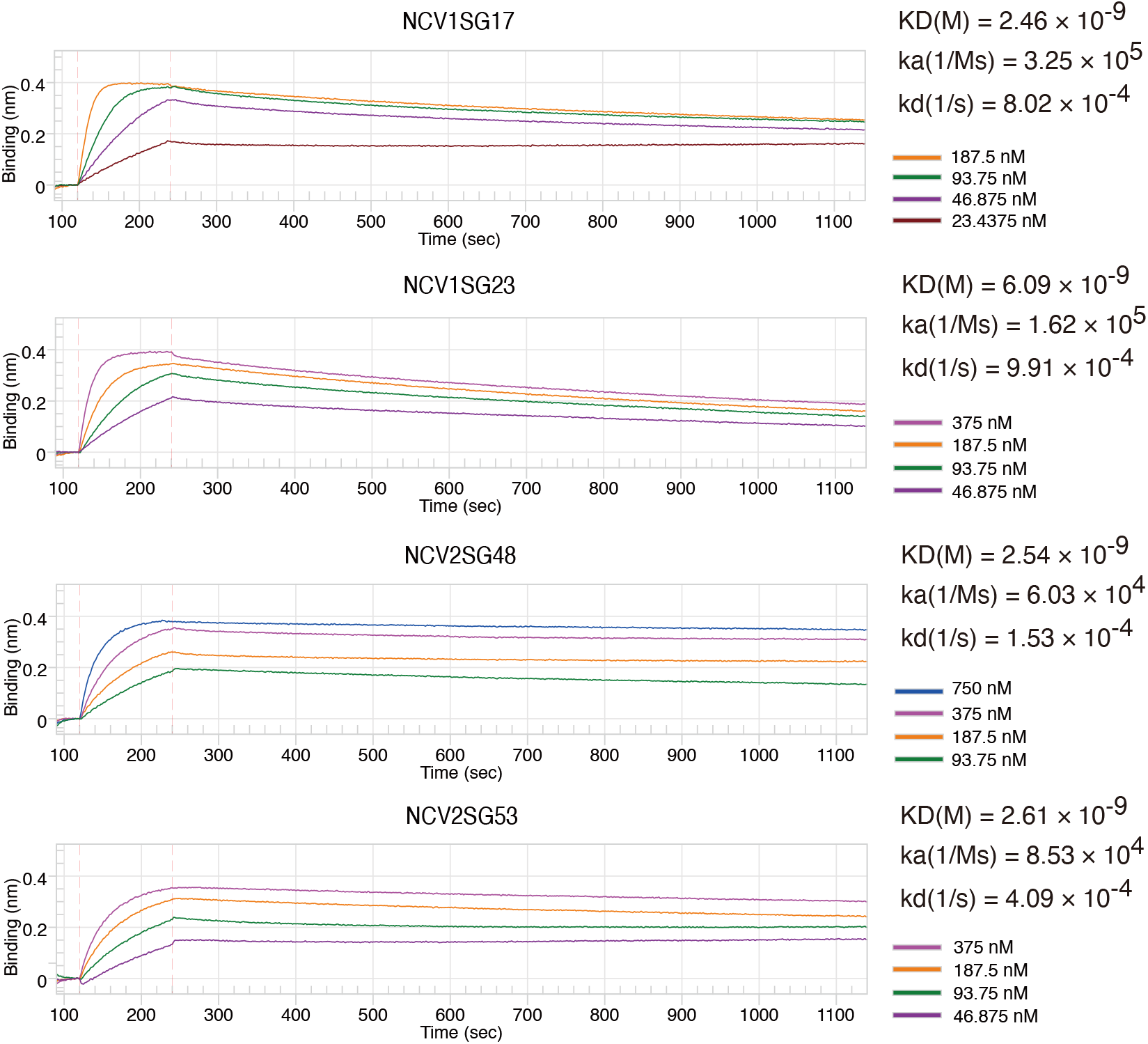
Binding affinity between Wuhan RBD and neutralizing mAbs. Biolayer interferometry results of NCV1SG17, NCV1SG23, NCV2SG48, and NCV2SG53 against the Wuhan RBD protein. Binding kinetics were measured for four different concentrations of the antigen and evaluated using a 1:1 binding model.

**Extended Data Fig. 4.**
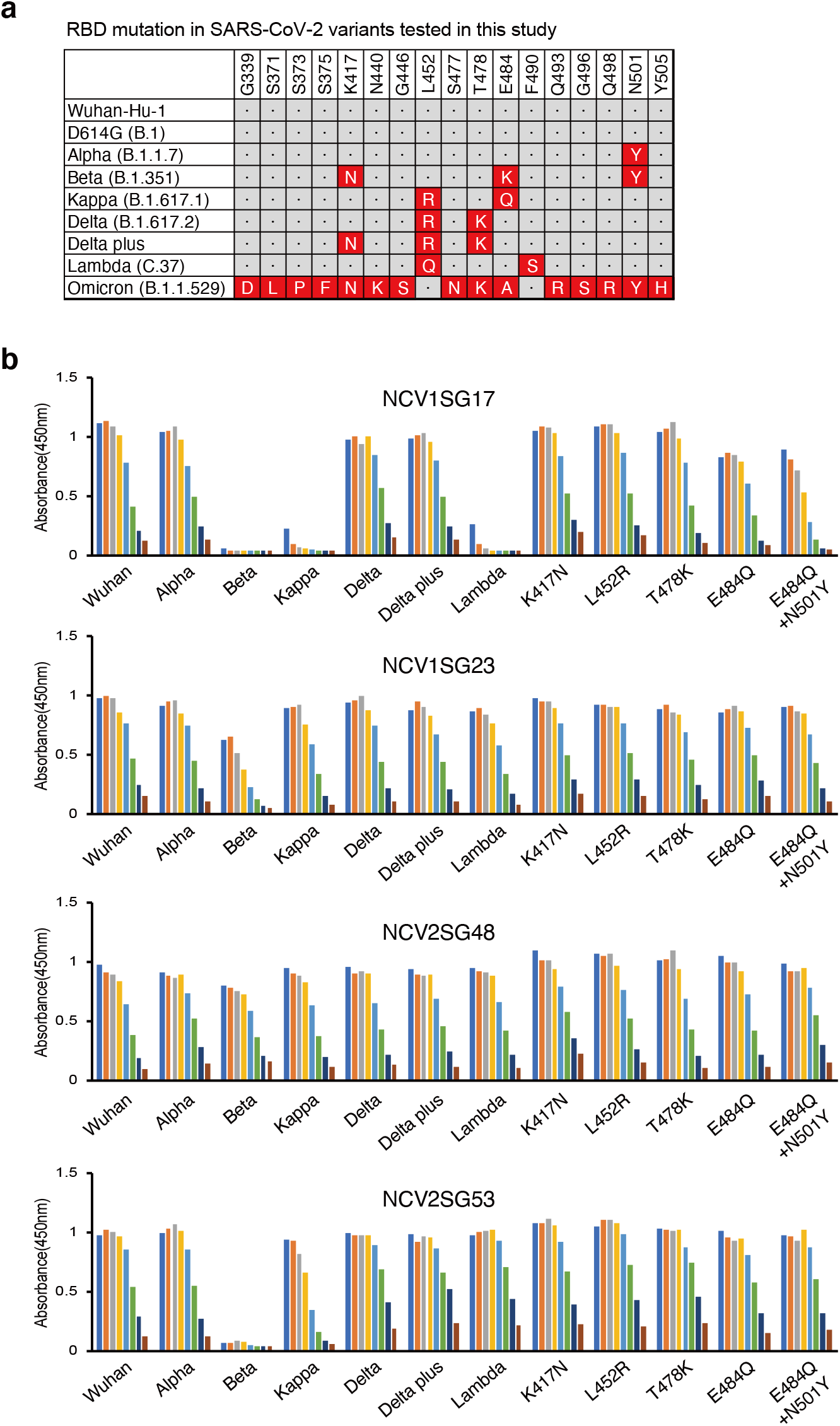
Binding of neutralizing mAbs to various SARS-CoV-2 RBD mutants. **a**, The matrix represents amino acid substitutions present in RBD of indicated SARS-CoV-2 variants. Name of variants are given on the *y-axis* and the position and amino acid replacement (single letter code) in each strain are given on the *x-axis*. **b**, The binding of neutralizing mAbs with S RBD protein of indicated variants or point mutants is determined by ELISA by three-fold serial dilutions.

**Extended Data Fig. 5.**
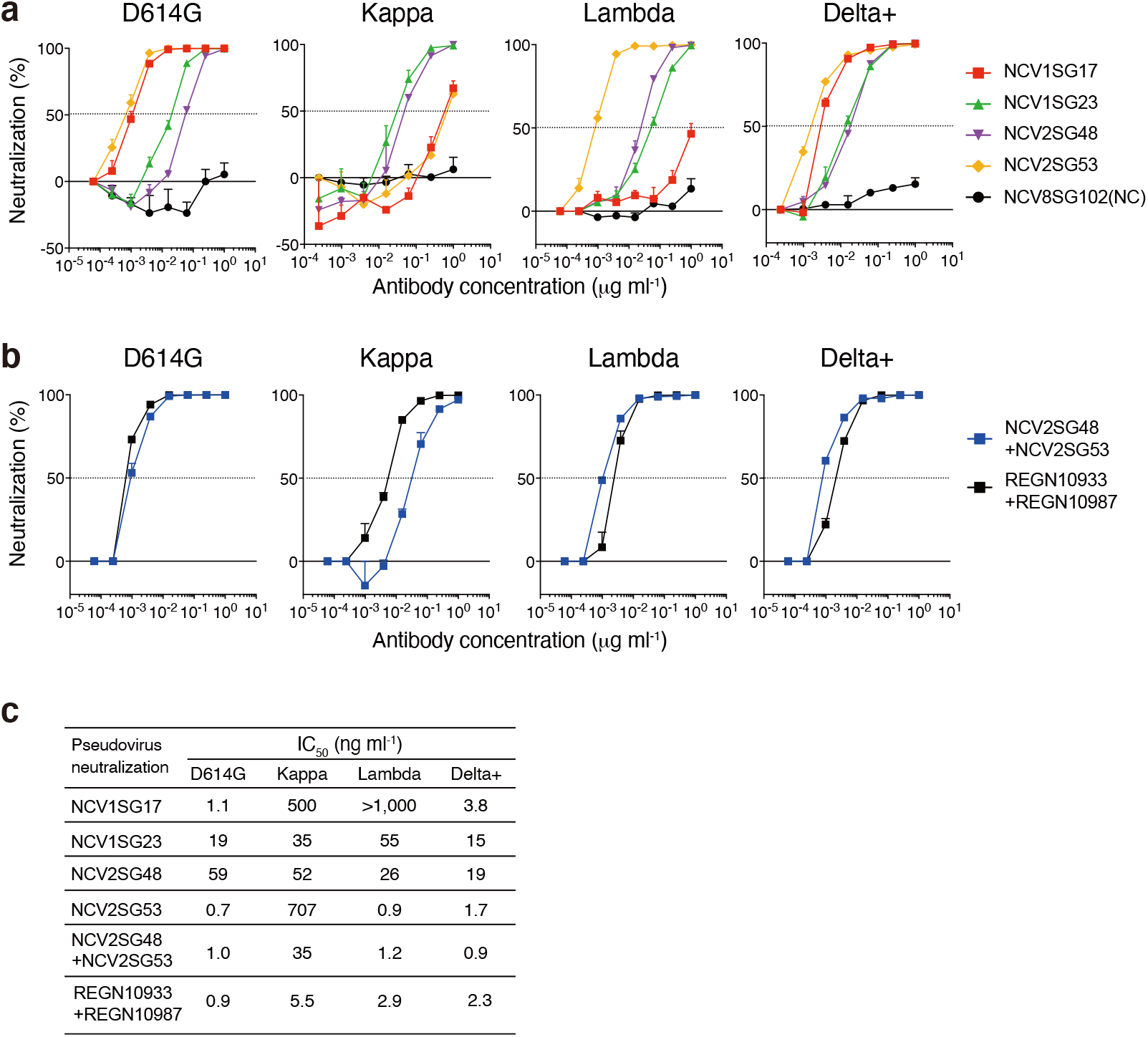
Neutralizing potency of mAbs against pseudoviruses. **a**, Dose-response analysis of the neutralization by each mAb NCV1SG17, NCV1SG23, NCV2SG48, NCV2SG53 on indicated variants of SARS-CoV-2 pseudovirus. The horizontal dotted line on each graph indicates 50% neutralization. Data are mean±SEM of technical duplicates from 2 to 3 independent experiments. **b**, Dose-response analysis of the neutralization by the mixture of mAbs NCV2SG48 and NCV2SG53, or REGN10933 and REGN10987 on indicated variants of SARS-CoV-2 pseudovirus. The horizontal dotted line on each graph indicates 50% neutralization. Data are mean±SEM of technical duplicates from 2 to 3 independent experiments. **c**, The IC_50_ of each antibody and the mixture of mAbs to indicated SARS-CoV-2 variants.

**Extended Data Fig. 6.**
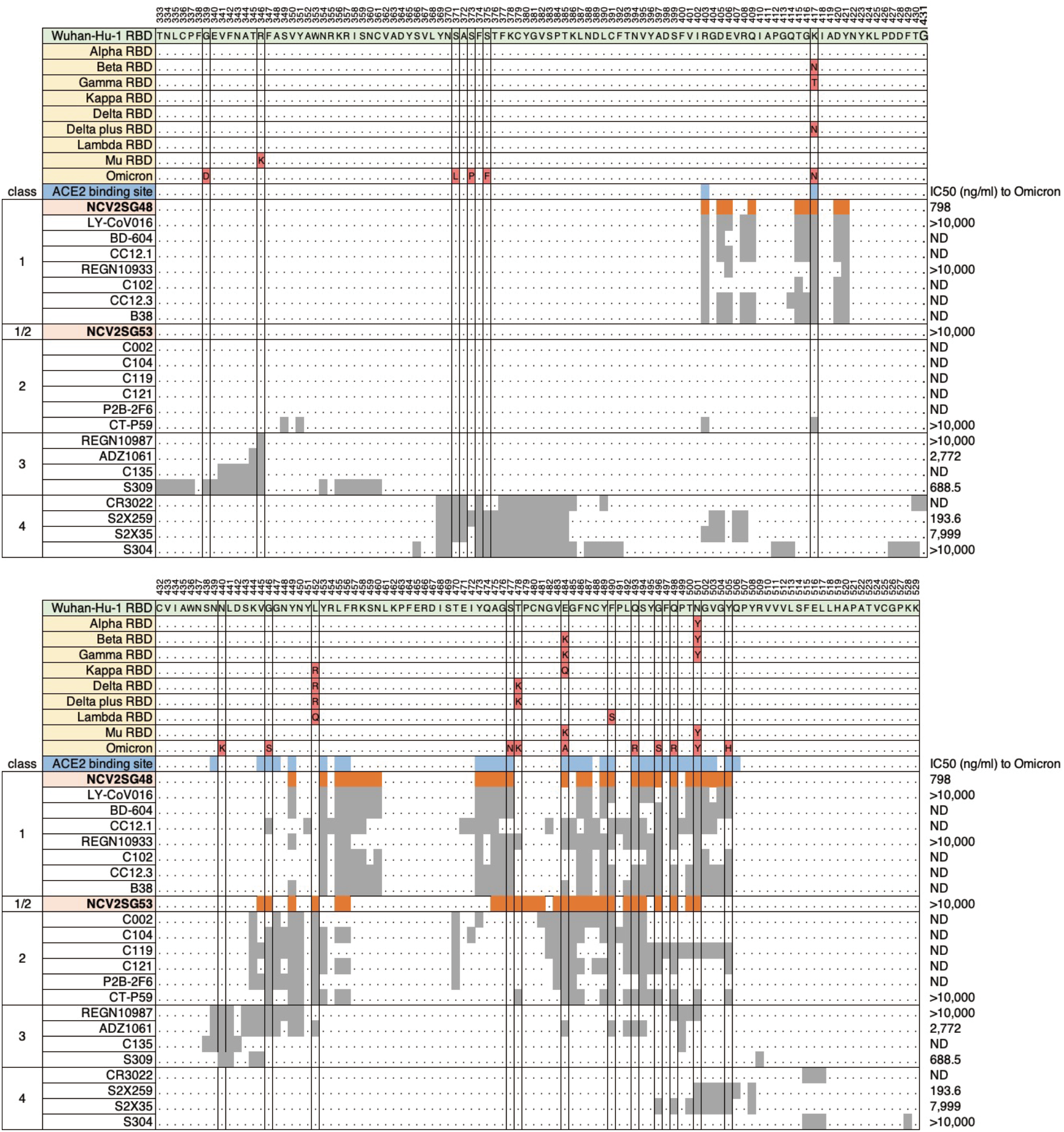
Mutations in SARS-CoV-2 RBD and epitope map of neutralizing antibodies on RBD sequence. RBD sequences, amino acid 333-431 (*top*) and 432-529 (*bottom*) of SARS-CoV-2 Wuhan-Hu-1 and variants are shown with replaced amino acids (*red*). ACE2 binding sites are shown in blue. The epitope of NCV2SG48 and NCV2SG53 determined by X-ray crystallography is shown in orange. Reported binding sites of class 1, class 2, class 3, and class 4 neutralizing mAbs are indicated in gray. We concluded NCV2SG48 as class 1 antibody and NCV2SG53 as class1/2 antibody. Amino acid substitutions acquired in variants are boxed. IC_50_ values against Omicron are shown on the right^6^.

**Extended Data Fig. 7.**
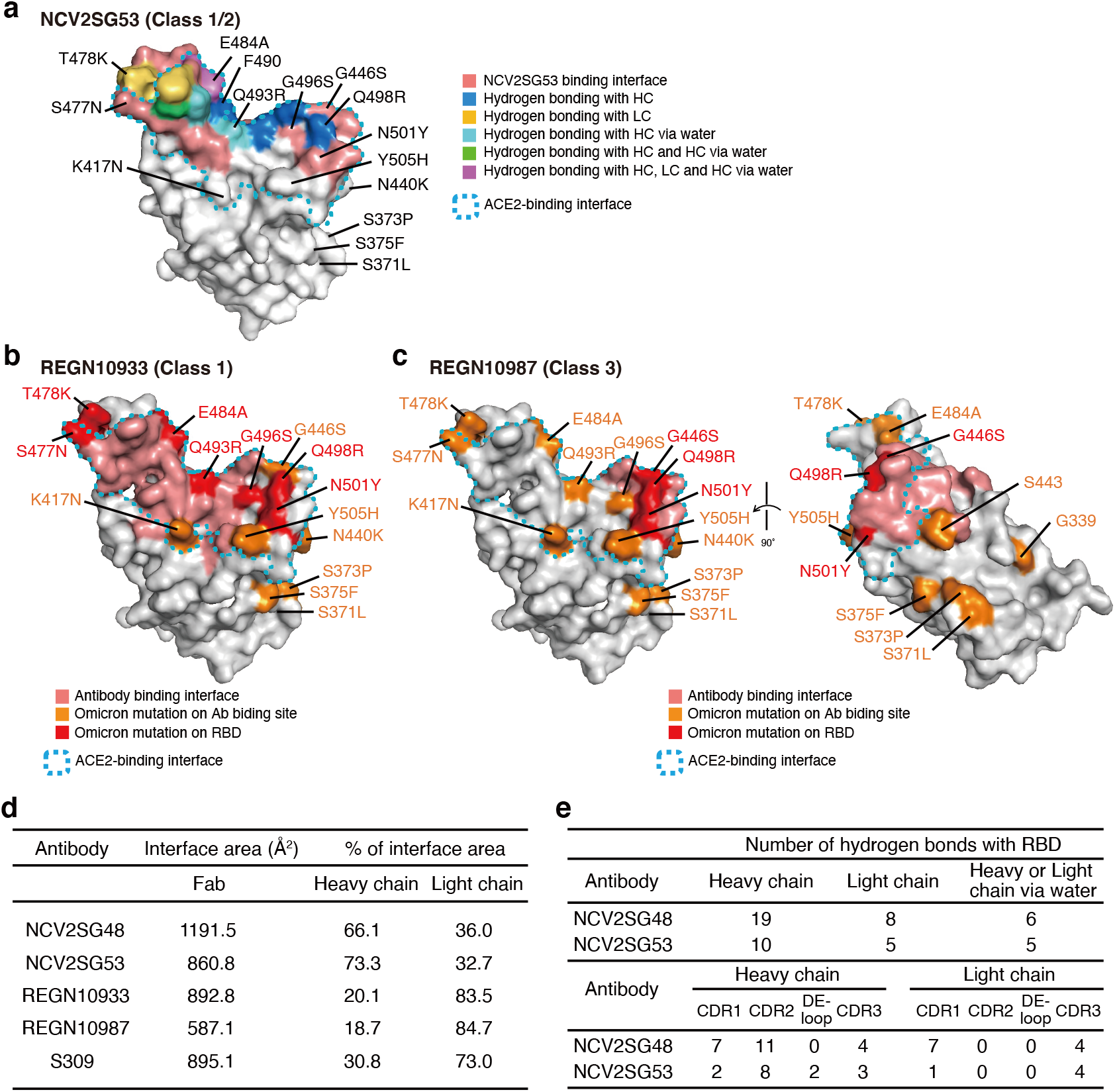
The structure and binding interface of a neutralizing antibody on the RBD indicate that NCV2SG48 has a large and universal interface than other mAbs. **a**, Antibody (NCV2SG53) binding sites are shown on the surface model of RBD. NCV2SG53 binding interface is shown in *light red* and the ACE2-binding site is surrounded by a *cyan* dotted line. Residues of hydrogen bonded to HC (*blue*), LC (*yellow*), and HC via water molecule (*cyan*) are shown on the RBD surface model. Residues from two hydrogen bonds with Fab are shown in *green* and more than three hydrogen bonds are shown in *magenta*. **b, c**, The binding interface of REGN10933 and REGN10987 on RBD. Antibody binding interface and Omicron mutations are shown by indicated colors. ACE2-binding interface is circled (*cyan*). **d**, Interface area of Fab from neutralizing antibody with RBD is calculated with percent of interface area of heavy chain and light chain. **e**, Number of hydrogen bonds between NCV2SG48 or NCV2SG53 and RBD.

**Extended Data Fig. 8.**
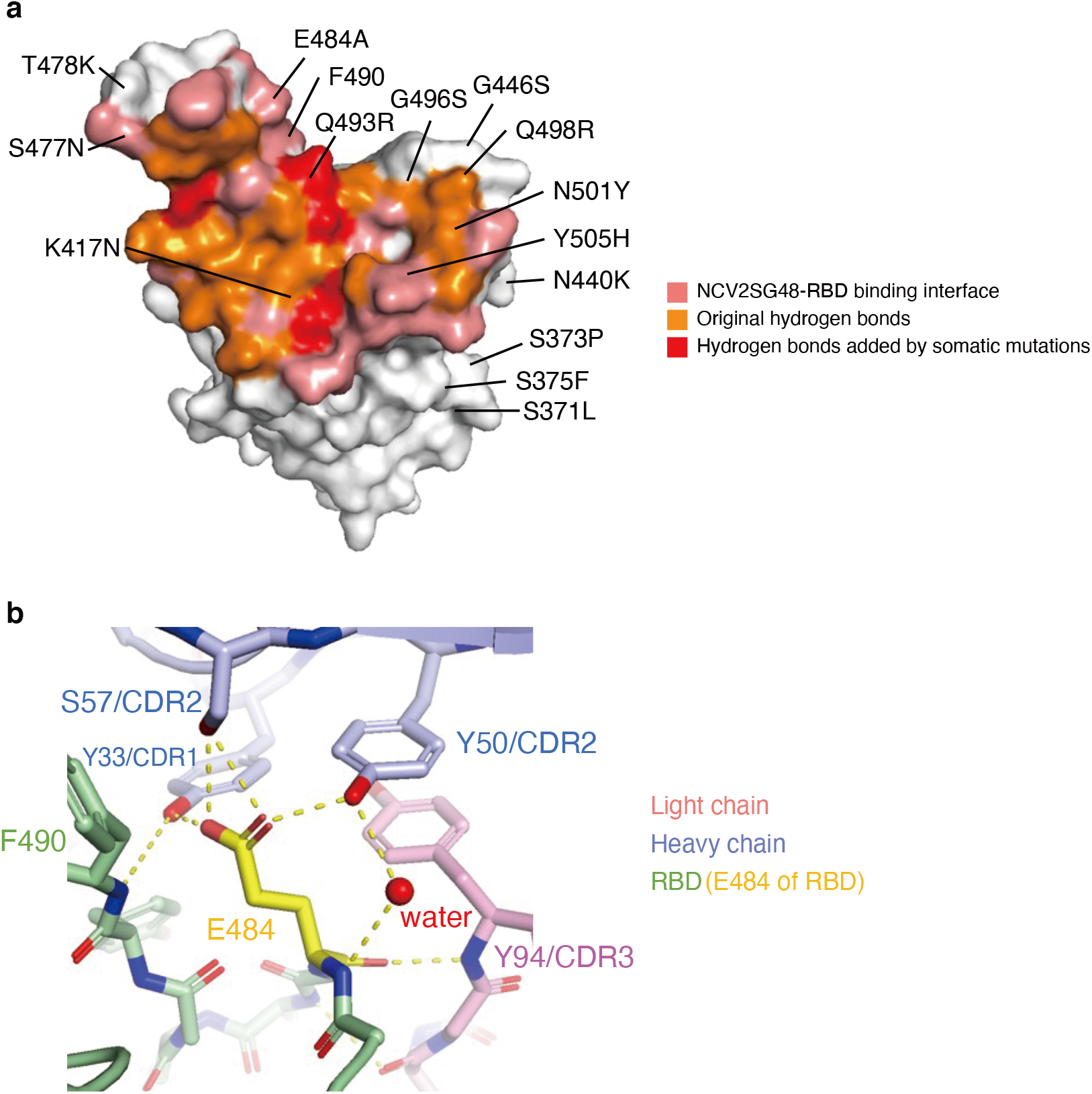
Somatic mutation contributes to hydrogen bonds between neutralizing mAbs and RBD. **a**, NCV2SG48-RBD binding interface. Somatic mutation contributed to creating additional hydrogen bonds for additional and stable interaction. Original hydrogen bonds and additional hydrogen bonds by somatic mutation are shown by red and orange, respectively. **b**, Interaction of RBD E484 residue with NCV2SG53 mAb through the multiple hydrogen bonds. Among hydrogen bonds formed on RBD E484 residue, S57 residue in the NCV2SG53 heavy chain CDR2 was generated by the somatic mutation and contributes to increased binding affinity.

**Extended Data Table 1.** Information of serum from vaccinated or convalescent individuals used in this study.

Vaccines
|  | Pfizer<br>(n=8) | Moderna<br>(n=6) | Total<br>(n=14) |
| --- | --- | --- | --- |
| Sex |  |  |  |
| Female | 6 | 2 | 8 |
| Male | 2 | 4 | 6 |
| Age (median, range) | 47 (31-62) | 38 (21-47) | 41 (21-62) |
| 1st shot | Mar 19-Jun 27, 2021 | Jul 5-6, 2021 | Mar 19-Jul 6, 2021 |
| 2nd shot | Apr 9-Jul 18, 2021 | Aug 5-6, 2021 | Apr 9-Aug 6, 2021 |
| Sampling days post-2nd dose; M 0.5 (median; range) | 15 (14-17) | 17 (16-18) | 16 (14-18) |
| Sampling days post-2nd dose; M 2 (median; range) | 66 (62-73) | 56 (56-57) | 62 (56-73) |
| Sampling days post-2nd dose; M5 (median; range) | 153 (152-163) | 152 (152) | 152 (152-163) |

Convalescents
|  | 17-55<br>(n=5) | 11-15<br>(n=6) | 8-10<br>(n=6) | Total<br>(n=17) |
| --- | --- | --- | --- | --- |
| Day from PCR+ |  |  |  |  |
| Sex |  |  |  |  |
| Female | 3 | 4 | 4 | 11 |
| Male | 2 | 2 | 2 | 6 |
| Age (median, range) | 71 (58-93) | 71 (59-87) | 55 (23-62) | 60 (36-93) |
| Date of PCR+ | Apr 14, 2020-<br>Dec 3, 2020 | Dec 8, 2020-<br>Jan 12, 2021 | Aug 11, 2020-<br>Dec 20, 2020 | Apr 14, 2020-<br>Jan 12, 2021 |
| Date of blood collection | Jun 8, 2020-Dec 20, 2020 | Dec 21, 2020-Jan 23, 2021 | Aug 20, 2020-Dec 28, 2020 | Jun 8, 2020-Jan 23, 2021 |

Demographics and vaccination information for serum samples from vaccinated individuals (*Vaccines*) and schedule of blood collection from COVID-19 convalescent individuals for serum and monoclonal antibody production (*Convalescents*).

**Extended Data Table 2.**
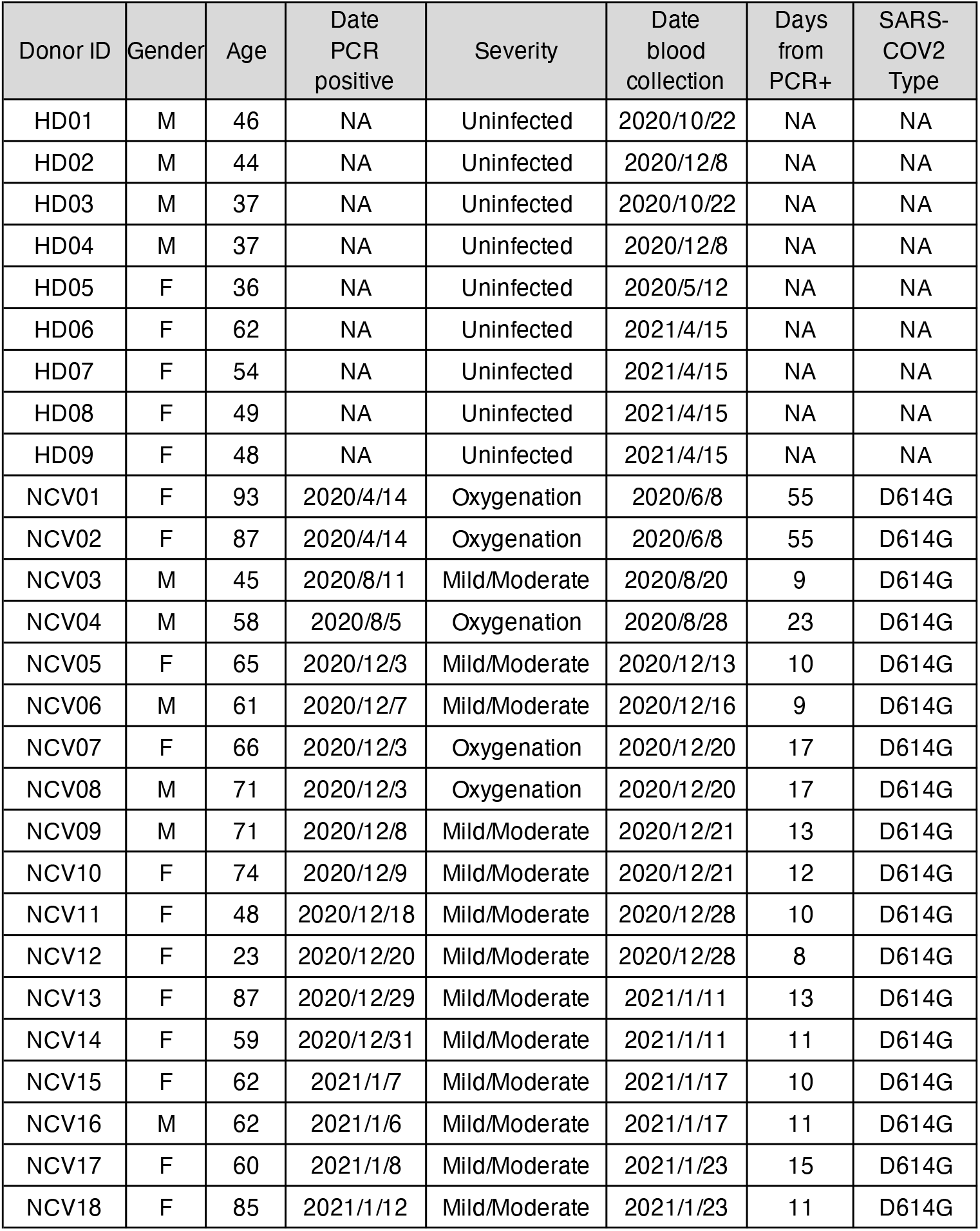
COVID-19 convalescent blood donor information used in this study.

**Extended Data Table 3.** Screening of escape mutants and neutralization results. **a**, Escape mutant screening was performed under the presence of NCV1SG17, NCV1SG23, NCV2SG48, or NCV2SG53 neutralization mAbs. Independent screening was repeated four to seven times for each mAb. **b**, Isolated escape mutants were sequenced to identify specific mutations and used for neutralization assay with a panel of neutralizing antibodies. MIC: minimum inhibitory concentration.

| **a** |  |  | **b** |  |  |  |  |  |
| --- | --- | --- | --- | --- | --- | --- | --- | --- |
| Antibody | SARS-CoV-2 virus (pfu) | Number of escape clone | Escape clone | Identified mutation | Neutralization, MIC (μg/ml) |  |  |  |
|  |  |  |  |  | NCV1 SG17 | NCV1 SG23 | NCV2 SG48 | NCV2 SG53 |
| NCV1SG17 | 2.2 x 10<sup>4</sup> | 0 | EM-17-1 | S494P | > 100 | < 2.5 | < 2.5 | 50-100 |
|  | 1.2 x 10<sup>5</sup> | 1 (EM-17-1) | EM-17-2 | S494P | > 100 | < 2.5 | < 2.5 | 50-100 |
|  | 5.6 x 10<sup>4</sup> | 4 (EM-17-2) | EM-17-3 | S494P | > 100 | < 2.5 | < 2.5 | 50-100 |
|  | 1.4 x 10<sup>5</sup> | 1 (EM-17-3) |  |  |  |  |  |  |
| Total | 3.4 x 10<sup>5</sup> | 6 |  |  |  |  |  |  |
| NCV1SG23 | 2.2 x 10<sup>2</sup> | 0 | EM-53-1 | E484D | > 100 | < 2.5 | < 2.5 | > 100 |
|  | 1.2 x 10<sup>3</sup> | 0 | EM-53-2 | G485D | > 100 | < 2.5 | < 2.5 | > 100 |
|  | 5.6 x 10<sup>2</sup> | 0 | EM-53-4 | G485R | > 100 | < 2.5 | < 2.5 | > 100 |
|  | 3.7 x 10<sup>3</sup> | 0 |  |  |  |  |  |  |
|  | 3.1 x 10<sup>4</sup> | 0 |  |  |  |  |  |  |
|  | 1.3 x 10<sup>4</sup> | 0 |  |  |  |  |  |  |
|  | 1.5 x 10<sup>4</sup> | 0 |  |  |  |  |  |  |
| Total | 6.4 x 10<sup>4</sup> | 0 |  |  |  |  |  |  |
| NCV2SG48 | 2.2 x 10<sup>2</sup> | 0 |  |  |  |  |  |  |
|  | 1.2 x 10<sup>3</sup> | 0 |  |  |  |  |  |  |
|  | 5.6 x 10<sup>2</sup> | 0 |  |  |  |  |  |  |
|  | 7.9 x 10<sup>2</sup> | 0 |  |  |  |  |  |  |
|  | 2.6 x 10<sup>4</sup> | 0 |  |  |  |  |  |  |
|  | 4.3 x 10<sup>4</sup> | 0 |  |  |  |  |  |  |
|  | 5.1 x 10<sup>4</sup> | 0 |  |  |  |  |  |  |
| Total | 1.2 x 10<sup>5</sup> | 0 |  |  |  |  |  |  |
| NCV2SG53 | 2.2 x 10<sup>3</sup> | 0 |  |  |  |  |  |  |
|  | 1.2 x 10<sup>4</sup> | 0 |  |  |  |  |  |  |
|  | 5.6 x 10<sup>3</sup> | 0 |  |  |  |  |  |  |
|  | 2.8 x 10<sup>4</sup> | 0 |  |  |  |  |  |  |
|  | 9.8 x 10<sup>4</sup> | 1 (EM53-1) |  |  |  |  |  |  |
|  | 1.1 x 10<sup>5</sup> | 1 (EM53-2) |  |  |  |  |  |  |
|  | 1.4 x 10<sup>5</sup> | 3 (EM53-4) |  |  |  |  |  |  |
| Total | 3.6 x 10<sup>5</sup> | 5 |  |  |  |  |  |  |
|  |  |  | Parental SARS-CoV-2 |  | < 2.5 | < 2.5 | < 2.5 | < 2.5 |

**Extended Data Table 4.**
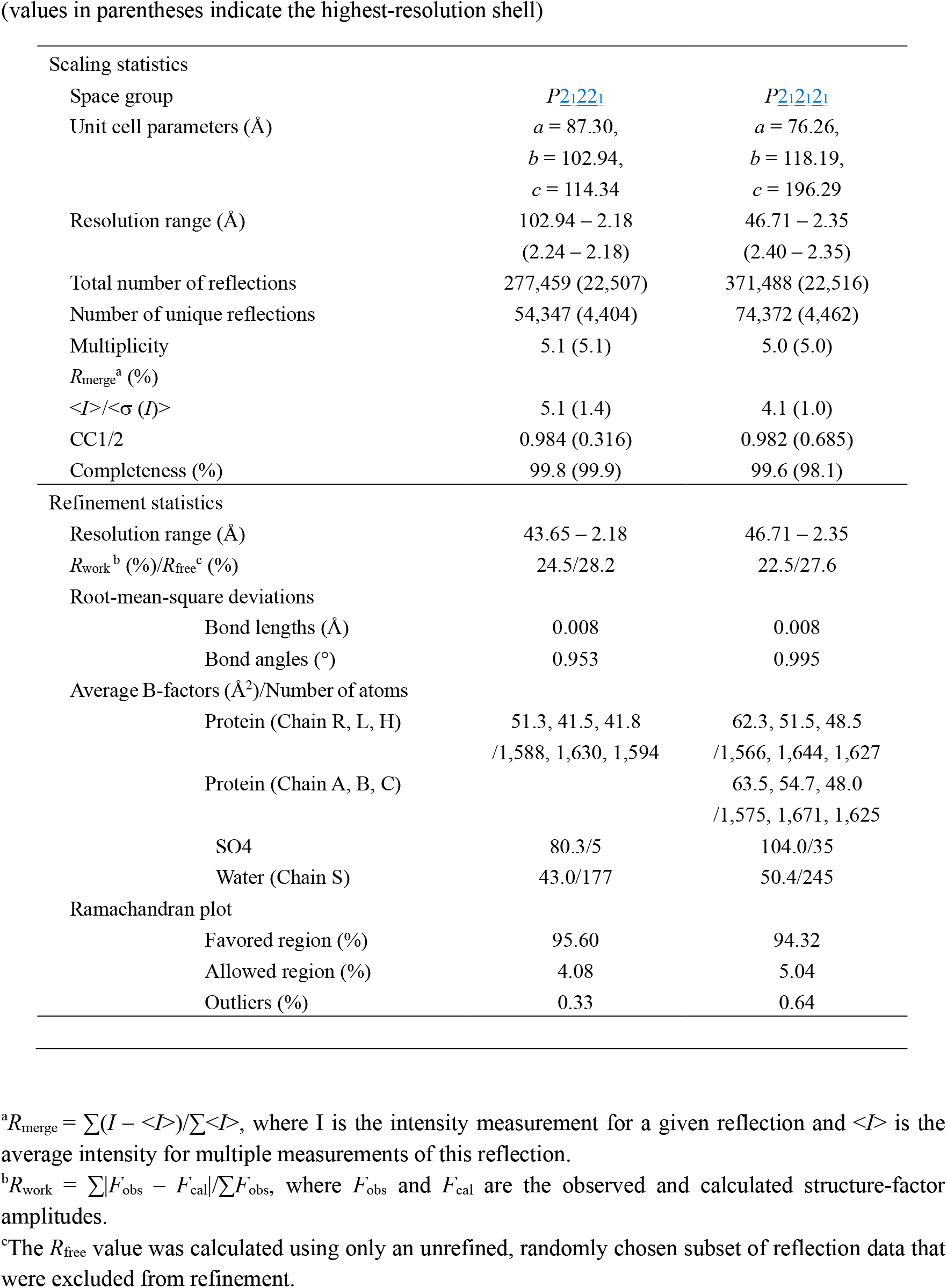
Data collection and refinement statistics.

